# FTY720 requires vitamin B_12_-TCN2-CD320 signaling in astrocytes to reduce disease in an animal model of multiple sclerosis

**DOI:** 10.1101/2022.01.10.475450

**Authors:** Deepa Jonnalagadda, Yasuyuki Kihara, Aran Groves, Manisha Ray, Arjun Saha, Hyeon-Cheol Lee-Okada, Tomomi Furihata, Takehiko Yokomizo, Edward V. Quadros, Richard Rivera, Jerold Chun

**Affiliations:** Sanford Burnham Prebys Medical Discovery Institute, 10901 N. Torrey Pines Rd, La Jolla, CA, 92037, USA; Neuroscience Graduate Program, School of Medicine, University of California San Diego, 9500 Gilman Dr, La Jolla, CA, 92093, USA; Department of Chemistry, University of Southern California, Los Angeles, California, 90089, USA; Department of Biochemistry, Graduate School of Medicine, Juntendo University, Hongo 2-1-1, Bunkyo-ku, Tokyo, 113-8421, Japan; Laboratory of Clinical Pharmacy and Experimental Therapeutics, School of Pharmacy, Tokyo University of Pharmacy and Life Sciences, Tokyo, 192-0392, Japan; Department of Medicine, SUNY-Downstate Medical Center, 450 Clarkson Ave, Brooklyn, NY 11203, USA

**Keywords:** Neuroinflammation, S1P_1_, FTY720, siponimod, ozanimod, ponesimod, lysophospholipid receptors

## Abstract

FTY720 (fingolimod) is a sphingosine 1-phosphate (S1P) receptor modulator and sphingosine analogue approved for multiple sclerosis (MS) therapy, which can functionally antagonize the S1P receptor, S1P_1_. Vitamin B_12_ (B_12_) deficiency produces neurological manifestations resembling MS. Here, we report a new mechanism where FTY720 suppresses neuroinflammation by regulating B_12_ metabolic pathways. Nuclear RNA-seq of c-Fos-activated astrocytes (called *ieAstrocytes*) from experimental autoimmune encephalomyelitis (EAE) spinal cords identified up-regulation of CD320, a transcobalamin 2 (TCN2)-B_12_ receptor, by S1P_1_ inhibition. CD320 was reduced in MS plaques. Deficiency of CD320 or dietary B_12_ worsened EAE and eliminated FTY720’s efficacy, while concomitantly down-regulating type I interferon signaling. TCN2 functioned as a chaperone for FTY720 and sphingosine, which induced astrocytic CD320 internalization. An accompanying paper identified a requirement for astrocyte sphingosine kinases in FTY720 efficacy and its altered expression in MS brains, molecularly linking MS and B_12_ deficiency that can be accessed by sphingolipid/fingolimod metabolic pathways.

## Introduction

Multiple sclerosis (MS) is a prototypical neuroinflammatory disease that produces demyelination and neurodegeneration in the CNS (Noseworthy et al., 2000). An animal model of MS – experimental autoimmune encephalomyelitis (EAE) – involves myelin antigen-primed helper T cells (Th1/Th17) and B cells (Gonzalez and Pacheco, 2014; Hauser et al., 2017) that attack the CNS to mimic MS. However, molecular mechanisms involving affected CNS cell types that can be therapeutically accessed to ameliorate disease in MS models and human MS remain incompletely understood. A long-recognized similar spectrum of neurological sequelae observed in MS also occurs with deficiency of vitamin B_12_ (B_12_; also called “cobalamin”) (Miller et al., 2005), suggesting a possibly overlapping disease mechanism; however, such linkage has been considered equivocal in the absence of molecularly defined pathways linking the two (Najafi et al., 2012). The B_12_ pathway involves formation of a complex between extracellular B_12_ and transcobalamin 2 (TCN2), which binds to multiple members of the low-density lipoprotein receptor (LDLR) family – megalin/LRP2, cubilin, and CD320/TCblR (Kozyraki and Cases, 2013) – enabling B_12_ to enter cells. In particular, CD320 mediates CNS B_12_ access (Alam et al., 2016).

Therapeutic efficacy in MS (and EAE) affecting both immune and CNS cell types has been realized through sphingosine 1-phosphate (S1P) receptor modulators (Chun et al., 2021; Chun et al., 2019). S1P is a lysophospholipid whose effects are meditated by cognate G protein-coupled receptors (GPCRs) (Mizuno and Kihara, 2020). FTY720 (known by its generic clinical name, fingolimod, which is a structural analogue of sphingosine) was the first FDA-approved orally available disease modifying therapy (DMT) for treating relapsing-remitting MS (RRMS) (Chun and Hartung, 2010; Chun et al., 2019; Cohen and Chun, 2011). FTY720 is phosphorylated by sphingosine kinases (SK1/2; gene name, *SPHK1/2*) to produce the active S1P analogue, FTY720P (fingolimod-P) that binds to four of the five known S1P GPCRs: S1P_1, 3, 4, 5_ (Brinkmann et al., 2002; Kihara et al., 2014; Kihara et al., 2015; Mandala et al., 2002; Mizuno and Kihara, 2020). In particular, the receptor subtype S1P_1_ is thought to be mechanistically important in MS through functional antagonism of S1P_1_ on lymphocytes, which reduces egress from secondary lymphoid organs (Arnon et al., 2011; Mandala et al., 2002; Matloubian et al., 2004). This has been proposed as a mechanism of action (MOA) of fingolimod by reducing pathogenic lymphocytes from entering the CNS to ameliorate disease (Chun et al., 2019).

However, direct CNS activities of S1P receptor modulators through expressed S1P receptors have also been proposed to occur in parallel with immunological activities (Chun and Brinkmann, 2011; Chun and Hartung, 2010; Cohen and Chun, 2011; Groves et al., 2013; Soliven et al., 2011), in view of the expression of S1P receptors in brain (Chun, 1999; Chun et al., 2000; Ishii et al., 2001; McGiffert et al., 2002; Zhang et al., 1999) and fìngolimod’s preferential accumulation within the CNS (Foster et al., 2007). In particular, EAE studies that employed astrocyte-specific S1P_1_ conditional knockout mice (S1P_1_-AsCKO) implicated S1P_1_ expressed on astrocytes for FTY720 efficacy (Choi et al., 2011; Groves et al., 2018). Conditional deletion of astrocyte S1P_1_ resulted in a loss of FTY720 efficacy despite the maintenance of immunological trafficking effects (Choi et al., 2011), supporting a distinct CNS mechanism which was consistent with fingolimod clinical data that reported reductions in brain volume loss in MS (Cohen and Chun, 2011; Kappos et al., 2006) and contrasting with increased brain volume loss produced primarily by anti-inflammatory agents (Vidal-Jordana et al., 2013).

To better understand the CNS cellular targets involved in EAE and MS, we previously used an unbiased *in vivo* screen to identify the earliest and most affected CNS cell types (Groves et al., 2018), as detected by immediate early gene *c-fos* expression, using a tetracycline transactivator (tTA)-controlled genetic tagging of c-Fos-activated cells that historically marked cellular activation with nuclear green fluorescence protein (GFP) (Matsuo et al., 2008). This unbiased screen identified “immediate-early astrocytes (*ieAstrocytes*),” accounting for over 95% of c-Fos-activated cells during EAE, whose prevalence linearly increased with EAE disease severity (Groves et al., 2018) consistent with *ieAstrocytes* contributing to the pathogenesis and progression of EAE. Moreover, both genetic removal or pharmacological inhibition of S1P_1_ suppressed *ieAstrocyte* formation (Groves et al., 2018), indicating that S1P-S1P_1_ signaling in *ieAstrocytes* has pivotal roles in FTY720 efficacy.

Here, the CNS molecular pathways accessed by FTY720 through astrocytes were investigated by analyzing gene expression profiles of isolated *ieAstrocytes*, under conditions of S1P_1_ inhibition, using fluorescence-activated nuclear sorting (FANS) combined with RNA-seq (Lake et al., 2016). This strategy identified a B_12_ pathway that was functionally validated using EAE and human MS brain samples, and further supported by an accompanying human brain MS resource dataset (Kihara, 2021). It also identified a second well-known type-I interferon (IFN-I) pathway relevant to MS. Notably, physical binding of FTY720 to TCN2, and CNS B_12_ signaling, provide molecular metabolic links to the long-recognized similarities between MS and B_12_ deficiency.

## Results

### *CD320 was identified by nuclear RNA-seq of* ieAstrocytes *and was up-regulated by S1P_1_ inhibition*

A nuclear RNA-seq pipeline developed for neurons (Lake et al., 2016) was adapted to assess *ieAstrocyte* nuclei isolated by FANS from EAE-induced WT^fos^, S1P_1_-AsCKO^fos^ (TetTag:S1P_1_^flox/flox^:hGFAP-cre), and WT^fos^+FTY720 mice (**Fig. 1A-C**). FANS enabled a more accurate assessment of activated astrocyte transcriptomes by focusing on GFP^+^ *ieAstrocytic* nuclei while additionally removing neurons by gating out NeuN^+^ nuclei (**Fig. 1B**). Both intronic and exonic reads were assessed from nuclear RNA which covered ~80% of transcripts in all three groups (**Fig. S1**). Control assessments of *S1pr1* (the mouse gene name for S1P_1_) expression revealed significant loss only in the S1P_1_-AsCKO^fos^ but not in WT^fos^ or WT^fos^+FTY720 nuclei (**Fig. S1**), supporting the expected gene deletion in astrocytes by the hGFAP promoter-driven Cre recombinase (Tien et al., 2012). In accordance with our previous findings and the definition of *ieAstrocytes* (Groves et al., 2018), *Fos* mRNA expression was decreased in both S1P_1_-AsCKO^fos^ and WT^fos^+FTY720 (**Fig. S1**). These nuclear RNA-seq datasets showed ~92% overlap with a reported gene set for mouse A1/A2 reactive astrocytes, but which lacked skewing towards either subtype (Liddelow et al., 2017) (**Fig. S1**), supporting a reactive yet distinct phenotype for *ieAstrocytes*.

**Fig. 1.**
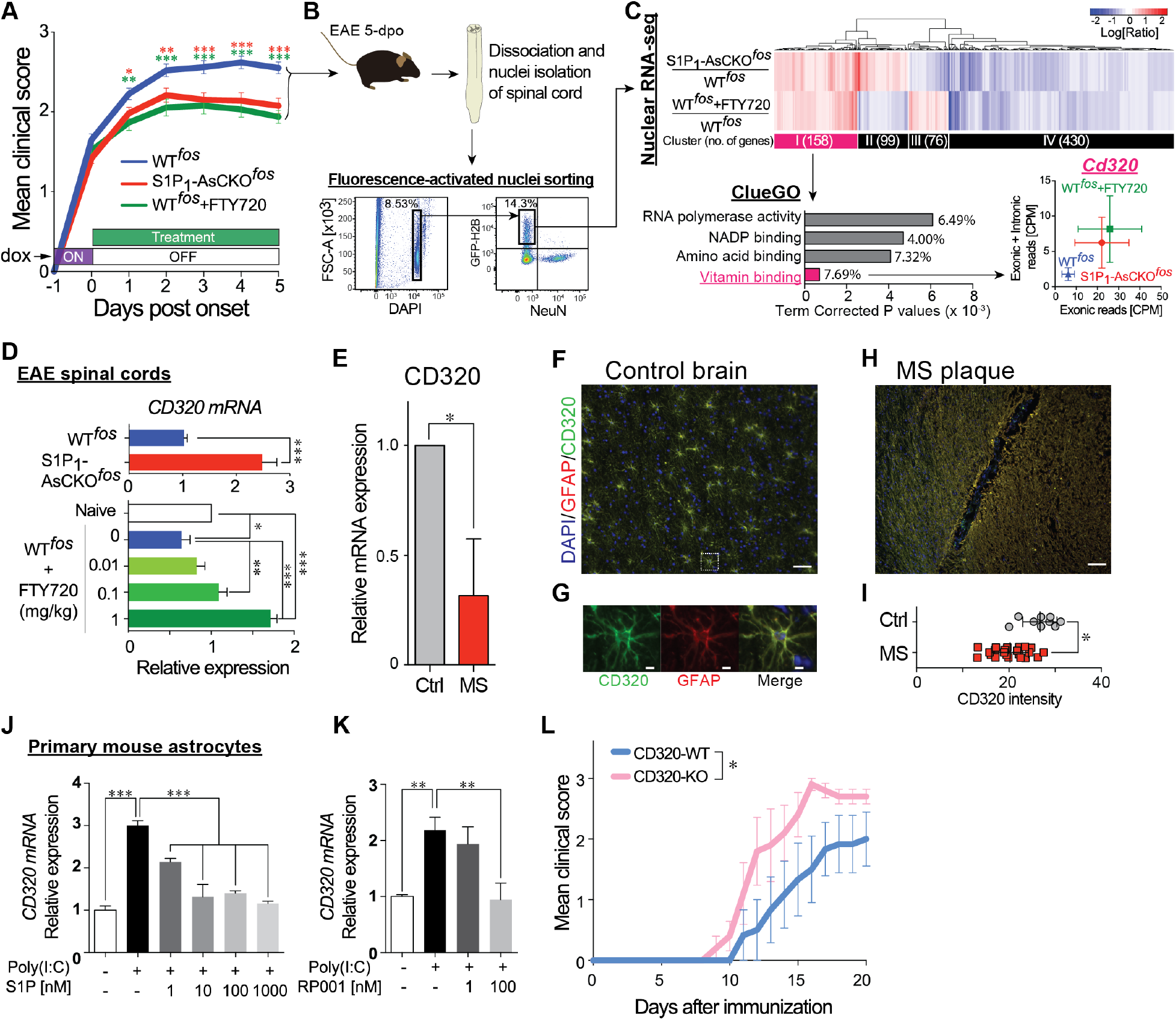
Identification of astrocyte CD320-vitamin B12 through studies of EAE and human multiple sclerosis brain supports CD320 loss in promoting neuroinflammation. **A**, Disease course of EAE-induced WT^fos^, S1P_1_-AsCKO^fos^, and FTY720-treated WT^fos^ (WT^fos^+FTY720) mice (n = 50, 43, and 32 animals, respectively). Day 0 was designated as the day of clinical sign onset (mean ± SEM, * p < 0.05; ** p < 0.01; *** p < 0.001 by two-way ANOVA with Bonferroni’s multiple comparisons test). The x-axis represents “days post onset,” which is different from “days after immunization.” **B**, Fluorescence-activated nuclei sorting (FANS) to isolate NeuN-GFP^+^ singlet nuclei from EAE spinal cords at 5-days post onset that is equivalent to 5-days off doxycycline. **C**, Nuclear RNA-seq. Heatmap is shown as 2-fold up- and down-regulated genes in S1P_1_-AsCKO^fos^ and WT^fos^+FTY720 vs. WT^fos^. Top 4 significantly enriched gene ontologies identified by ClueGO (Bindea et al., 2009) of Cytoscape (Cline et al., 2007) are shown. Expression levels of *Cd320* (count per million, CPM) are plotted. **D**, Expression of *Cd320* mRNA in EAE SCs determined by qPCR (mean ± SEM, * p < 0.05, ** p < 0.01 and *** p < 0.001 by t test for top panel, and one-way ANOVA with Tukey’s multiple comparisons test for bottom panel). **E**, *Cd320* mRNA expression in human MS normalized to control brains (mean ± SEM, n = 5 and 2 brains, respectively. * p < 0.01 by one sample t test). **F-I**, Immunohistochemistry (IHC) for CD320 and GFAP in human control brain (**F** and **G**) and MS brain disease plaques (**H**). Scale bar, 100 μm (**F** and **H**) and 5 μm (**G**). CD320 immunolabeling is significantly reduced in human MS plaques as compared to control brain (**I**). Each symbol represents an assessed region of interest in the sections. * p < 0.001 by Mann-Whitney U test. **J-K**, *Cd320* mRNA expression in primary astrocytes stimulated with 10 μg/mL poly(I:C) in the presence of S1P (**J**) or S1P_1_ specific agonist, RP001 (**K**). Data are from three independent experiments of three technical replicates (mean ± SEM, ** p < 0.01 and *** p < 0.001 by one-way ANOVA with Tukey’s multiple comparisons test). **L**, EAE disease course of CD320-WT and CD320-KO mice (n = 6 and 5 animals, respectively). Data are from three independent experiments (mean ± SEM, *, statistical significance was analyzed by two-way ANOVA; interaction, p = 0.90; time, p <0.0001; genotype, p < 0.001).

Hierarchical clustering of 2-fold up- or down-regulated genes in S1P_1_-AsCKO^fos^ and WT^fos^+FTY720 vs. WT^fos^ nuclei revealed four distinct clusters (I-IV) (**Fig. 1C** and **Table S1**). Cluster I consisted of 158 up-regulated genes that were of primary interest in identifying anti-neuroinflammatory genes produced by S1P_1_ inhibition. The ClueGO plugin (Bindea et al., 2009) of Cytoscape (Cline et al., 2007) identified a vitamin binding pathway as the most significantly enriched Gene Ontology (GO) molecular function term (**Fig. 1C**). Reactome pathway analysis (Fabregat et al., 2017) identified 27 pathways (p < 0.05, **Table S2**) composed of 8 genes (*Cd320, Dhfr, Tlr3, Eif2ak2, Ppp2cb, Hist1h2bf, Cdc7*, and *Psmb2*). Further analyses by quantitative PCR (qPCR) of these 8 genes in the spinal cords (SCs) of chronic EAE mice revealed that *Cd320* was significantly downregulated during EAE (**Fig. S1**), which was supported by previously reported RNA-seq data that compared naive vs. EAE astrocytes (~20%) (Rothhammer et al., 2016). Critically, *Cd320* expression was increased during amelioration of EAE through S1P_1_ genetic deletion or pharmacological inhibition by FTY720 exposure in animals (**Fig. 1D**).

### CD320 expression was down-regulated in human MS brains

CD320 is a Type I membrane protein that belongs to the LDLR family that transports the TCN2-B_12_ complex into CNS cells (Quadros and Sequeira, 2013). It has a primary role in maintaining brain B_12_ homeostasis, as demonstrated by greater than 95% reduction of B_12_ in the CNS of CD320-KO mice (Lai et al., 2013). To assess the human relevance of CD320 downregulation in MS, CD320 expression was assessed in human brains by qPCR, identifying significant reduction (~70%) in MS plaques as compared to non-diseased controls (**Fig. 1E**). These results are consistent with publicly available datasets derived from chronic MS plaques (Han et al., 2012). CD320 immunoreactivity was examined in human brain sections containing MS plaques as compared to non-diseased controls (**Fig. 1F-I**) and reduced CD320 immunoreactivity was identified on GFAP^+^ astrocytes in plaques compared to controls (**Fig. 1I**). These results support the relevance of CD320 and its related pathways to the human MS brain.

### *Analyses* in vitro *and* in vivo *supported worsening of disease with CD320 loss*

The restoration of *Cd320* expression during EAE by genetic or pharmacological S1P_1_ inhibition (**Fig. 1D**) indicated that S1P_1_ activation negatively regulates CD320 expression. Primary mouse astrocytes were examined in culture by first up-regulating (2~3-fold) *Cd320* expression via exposure to polyinosinic-polycytidylic acid (poly(I:C)), a Toll-like receptor 3 (TLR3) agonist (Alexopoulou et al., 2001) that was reported to ameliorate EAE (Khorooshi et al., 2015) and was thus anticipated to increase *Cd320* expression, which was observed (**Fig. 1J, K**). Both S1P and a short-acting S1P_1_-specific agonist, RP001 (Cahalan et al., 2011), inhibited poly(I:C)-induced *Cd320* expression in a dose-dependent manner (**Fig. 1J, K**) consistent with increased S1P levels reported in human MS brain lesions (Miller et al., 2017) and the reduced CD320 observed here (**Fig. 1E-I**). Linkage between S1P_1_ and CD320 predicted that genetic removal of CD320 should exacerbate EAE. To test this hypothesis, CD320-WT and CD320-KO mice were challenged with EAE, which resulted in a more severe disease course with earlier onset in the CD320-KO compared to WT controls (**Fig. 1L**, **Fig. S1**) that was also independent of peripheral blood lymphocyte counts (**Fig. S1**). These results showed that CD320 loss exacerbates EAE.

### B_12_ restriction also worsened EAE with parallel activation of inflammatory responses and suppression of IFN-I signaling

Down-regulation of CD320 in both EAE and MS may result in B_12_ deficiency within the CNS, although the effects of B_12_ deficiency on EAE severity have not been previously reported. B_12_ is an essential vitamin obtained through diet (Nielsen et al., 2012), and therefore, mice deficient in B_12_ were produced (B_12_^def^) through ~8-9 weeks of dietary restriction (Ghosh et al., 2016), followed by EAE challenge. B_12_^def^ mice exhibited severe EAE (**Fig. 2A, Fig. S2**), significant B_12_ reduction in SCs (**Fig. 2B**), increased histological damage (**Fig. 2C**), but equivalent T cell proliferative responses against MOG_35-55_ peptide as compared to controls (**Fig. S2**). In addition, CD320 protein levels were also down-regulated (by ~40%) in B_12_^def^-EAE SCs compared to controls (**Fig. S2**).

**Fig. 2.**
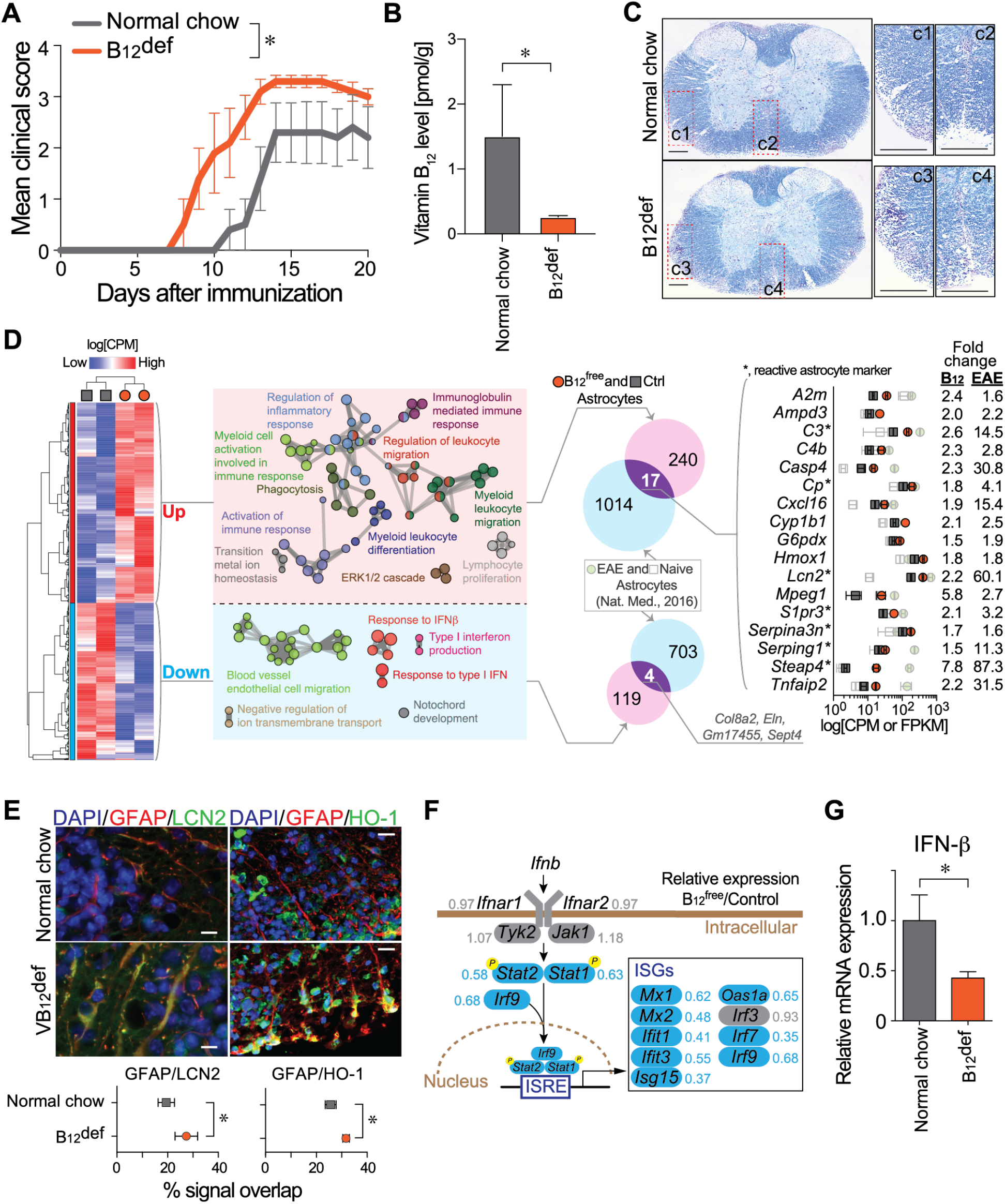
Vitamin B_12_ restriction worsens EAE, activates astrocytes, and reduces IFN-I signaling. **A**, EAE disease course of control (normal chow) and B_12_-deficient (B_12_^def^) diet fed mice (n = 5 animals). Data are from two independent experiments (mean ± SEM; *, statistical significance was analyzed by two-way ANOVA; interaction, p = 0.32; time, p <0.0001; diet, p < 0.001). **B,** Vitamin B_12_ levels in the EAE spinal cords (n = 6; mean ± SEM; *, p < 0.01 by Mann Whitney U test.) **C,** Luxol fast blue staining of EAE SCs at 15 days post immunization. Regions of interest (c1-c4) are magnified. Scale bar, 200 μm. **D**, Heatmap for up- and down-regulated differentially expressed genes (DEGs) produced from RNA-seq data of cultured astrocytes in serum-free B_12_^free^ and control media. DEGs were analyzed using ClueGO plugin (Bindea et al., 2009) of Cytoscape (Cline et al., 2007), and the functional groups shown as networks with different colors. DEGs were compared with EAE astrocytes (Rothhammer et al., 2016), which identified 17 and 4 commonly up- and down-regulated genes, respectively. Expression levels of commonly up-regulated genes in B_12_^free^/Ctrl [CPM] and EAE/Ctrl [FPKM; fragment per kilobase of transcripts per million] are plotted. **E**, IHC validation for up-regulated genes; LCN2 or HO-1 (green) and GFAP (red) in the EAE SC at 21 days post immunization. Scale bar, 10 μm. Overlapping signals of LCN2 or HO-1 in GFAP^+^ astrocytes are plotted (mean ± SEM, n = 5 animals, * p < 0.05 by unpaired t test). **F**, Detailed mapping of DEGs within the astrocyte signaling cascade for the IFN-I system. Gene names along with their respective ratio of B_12_^free^ to control are shown. ISRE, IFN-stimulated response element. ISGs, IFN-stimulated genes. **G**, IFN-β mRNA expression in the EAE SCs was determined by qPCR ( *, p < 0.05 by unpaired t test).

To explore the mechanisms underlying increased EAE severity in B_12_-restricted animals, the transcriptomic consequences of B_12_ loss in astrocytes were examined. Primary mouse astrocyte cultures were maintained for 2 weeks in B_12_-free (B_12_^free^) vs. control media, both of which supported growth and GFAP positivity. RNA-seq analysis identified 257 up-regulated and 123 down-regulated differentially expressed genes (DEGs; showing >1.5-fold, p < 0.05; or < 0.6-fold, p < 0.05, respectively: **Fig. 2D** and **Table S3**). Gene Ontology (GO) analyses of up-regulated DEGs revealed enrichment of multiple pro-inflammatory cascades (**Fig. 2D**). Although no obvious astrocyte class A1/A2-skewing (Liddelow et al., 2017) was observed (**Fig. S2**), B_12_^free^ astrocytes shared 17 DEGs with EAE astrocytes (Rothhammer et al., 2016) (**Fig. 2D**), in which 7 genes were shared with A1/A2 reactive astrocyte genes (Liddelow et al., 2017). Protein expression of up-regulated genes in astrocytes was supported by co-immunolabeling for GFAP and proteins for two identified genes, lipocalin-type 2 (LCN2/*Lcn2*) and heme oxygenase-1 (HO-1/*Hmox1*), in B_12_^def^ EAE SCs (**Fig. 2E**). Interestingly, despite no exposure to inflammatory stimuli, B_12_^free^ astrocytes displayed a reactive astrocyte phenotype, implicating B_12_-CD320 in maintaining astrocytes in quiescent or non-reactive states.

Further GO analyses revealed a significant enrichment of neuroprotective IFN-I signaling pathways whose genes were markedly downregulated in B_12_^free^ astrocytes as compared to controls (**Fig. 2F**). The mRNA expression of an endogenous ligand for IFN-I receptors – IFN-β – showed significant down-regulation in B_12_^def^ EAE SCs over controls (**Fig. 2G**), which appeared to be produced by microglia in the EAE-affected CNS (Khorooshi et al., 2015; Kocur et al., 2015). Taken together, loss of B_12_ promotes neuroinflammation by inducing astrocyte pro-inflammatory responses, reducing IFN-I sensitivity in astrocytes, and suppressing IFN-β production in microglia and possibly other immune cell types.

### Loss of CD320 or B_12_ eliminated efficacy of FTY720 in EAE

S1P_1_ inhibition restored CD320 expression in the CNS (**Fig. 1D**), raising the possibility that FTY720’s beneficial effects in EAE could be reduced or diminished in CD320-KO mice or dietarily B_12_-restricted mice. EAE-induced WT mice, which served as positive controls, exhibited the expected ameliorating effects of FTY720 treatment on EAE scores (**Fig. 3A**), recovery rate (**Fig. 3B**), expected reductions in peripheral blood lymphocyte counts (**Fig. S3**), and reduced IFN-β mRNA expression (**Fig. 3C**). In marked contrast, FTY720 did not ameliorate EAE in CD320-KO mice (**Fig. 3A, B**), yet still reduced peripheral blood lymphocyte numbers (**Fig. S3**). Moreover, FTY720 did not significantly restore IFN-β mRNA expression in CD320-KO mice (**Fig. 3C**).

**Fig. 3.**
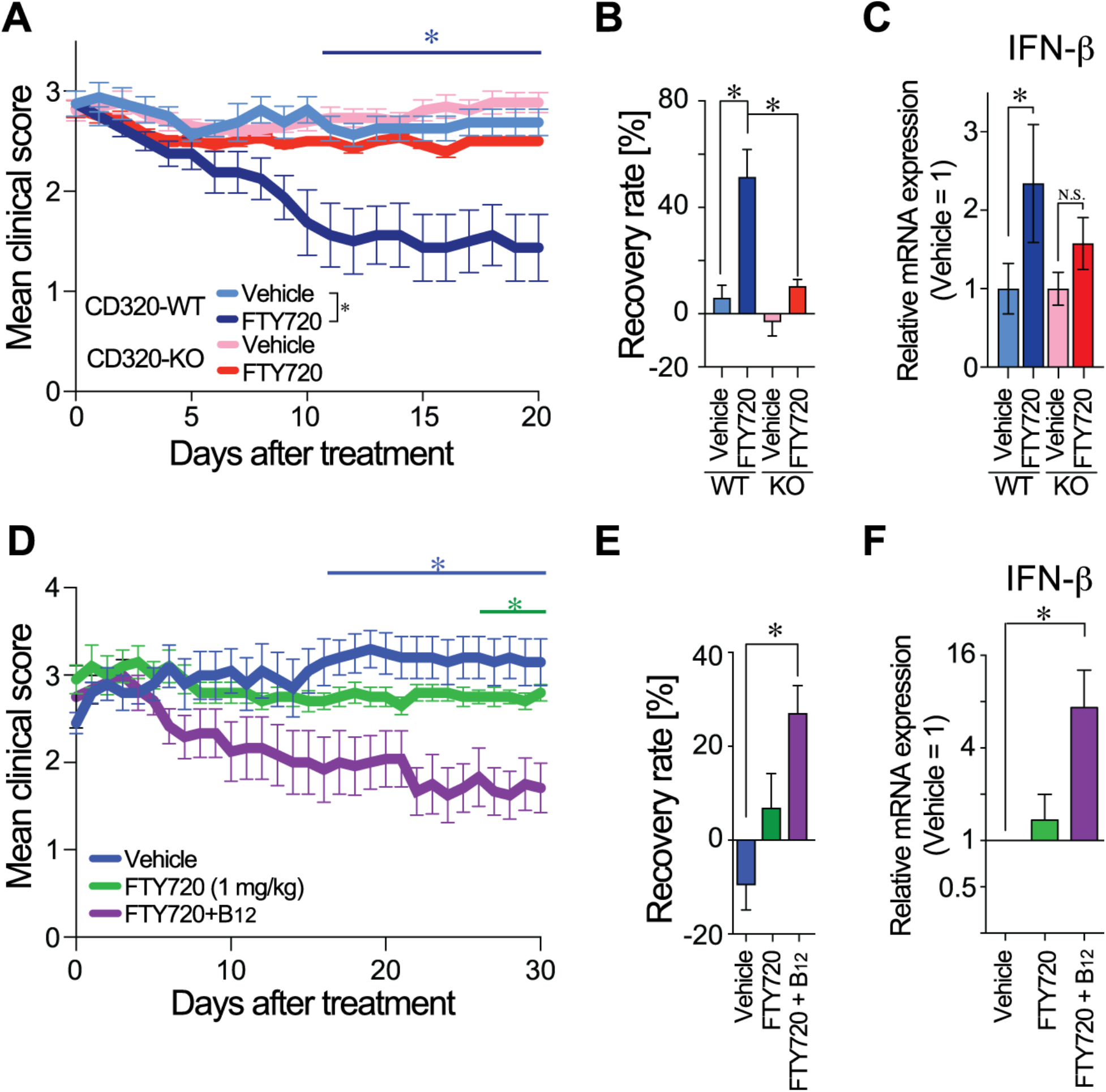
Loss of FTY720 efficacy during EAE in CD320-deficient and dietary B_12_-restricted mice. **A**, FTY720 treatment on EAE-induced CD320-WT (n = 8) and CD320-KO mice (n = 14). Data are combined from two independent experiments (mean ± SEM; *, statistical significance was analyzed by two-way ANOVA between day 9 to 20; interaction, p = 0.16; time, p <0.01; genotype, p < 0.01). **B,** Recovery rate with FTY720 treatment (*, p < 0.05 by one-way ANOVA with Bonferroni’s multiple comparisons test). **C,** IFN-β mRNA expression in EAE SCs (*, p < 0.05 by one-way ANOVA with Bonferroni’s multiple comparisons test; *N.S*., non-significant). **D,** Therapeutic treatment of EAE-induced B_12_^def^ mice with vehicle, FTY720, and FTY720+B_12_ (n = 5, 5, 6 animals, respectively). Data are representative of 2 independent experiments (mean ± SEM; * p < 0.05 vs. Vehicle by two-way ANOVA, interaction, p = 0.27, time, p = 0.74; diet, p < 0.0001 with Bonferroni’s multiple comparisons test). **E,** Recovery rate with FTY720 treatment (*, p < 0.05 by one-way ANOVA with Bonferroni’s multiple comparisons test). **F,** IFN-β mRNA expression in EAE SCs (*, p < 0.05 by one-way ANOVA with Bonferroni’s multiple comparisons test).

Since B_12_ is essential for protein synthesis, its deficiency suppressed CD320 expression (**Fig. S2**). B_12_^def^-EAE mice might also lose responsivity to FTY720. As was the case in CD320-KO mice, FTY720 lost efficacy in B_12_^def^-EAE mice (**Fig. 3D, E**), even though FTY720 reduced peripheral blood lymphocyte counts (**Fig. S3**). Importantly, B_12_^def^-EAE mice that were administered combination therapy (FTY720+B_12_) exhibited the greatest response in reducing disease score (**Fig. 3D**), an increased recovery rate (**Fig. 3E**), and elevated IFN-β mRNA expression in SCs (**Fig. 3F**). These responses correlated with the recovery of CD320 expression observed in the FTY720+B_12_ group (**Fig. S3**). Collectively, CD320 or B_12_ were independently required for FTY720 efficacy in EAE while producing synergistic therapeutic effects with B_12_ and FTY720.

### FTY720 bound to TCN2 with subnanomolar affinity and induced CD320 internalization

The loss of FTY720 efficacy in CD320-KO mice (**Fig. 3A**) generated a hypothesis that CD320 might be a carrier for FTY720 via direct binding. This was tested using a compensated interferometric reader (CIR) that is a label-free, optical technique for measuring molecular interactions in free solution (Mizuno et al., 2019; Ray et al., 2020). A positive control included CD320 binding to TCN2 (**Fig. 4A**), whose K_D_ value (1.86 nM; 95% CI, 0.13~17.05; R^2^ = 0.91) was comparable to previously published values (Alam et al., 2016). However, no binding signals were observed between CD320 and FTY720 or FTY720P (**Fig. 4B**). By comparison, sphingosine (Sph: K_D_ = 0.36 nM; 95% CI, −0.01~73.2; R^2^ = 0.81) and S1P (K_D_ = 11.8 nM; 95% CI, 0.56~infinity; R^2^ =0.91) did show binding to CD320 (**Fig. 4C**). Since the hypothesis of FTY720-CD320 interactions was rejected, an alternative hypothesis was that FTY720 instead directly binds to TCN2. As expected, TCN2 bound B_12_ (KD = 0.19 nM; 95% CI, −0.19~11.2; R2 = 0.83) (**Fig. 4D**). TCN2 also bound both FTY720 (K_D_ = 0.24 nM; 95% CI, 0.03~1.79; R^2^ = 0.96; **Fig. 4E**) and Sph (K_D_ = 0.14 nM; 95% CI, −0.02~49.6; R^2^ = 0.88; **Fig. 4 F**), whereas little or no binding was identified for FTY720P (**Fig. 4E**) and S1P (**Fig. 4 F**). A next generation S1PR modulator, BAF312 (siponimod) that is not a sphingosine analogue and does not require *in vivo* phosphorylation, did not show binding to TCN2 (**Fig. S4**).

**Fig. 4.**
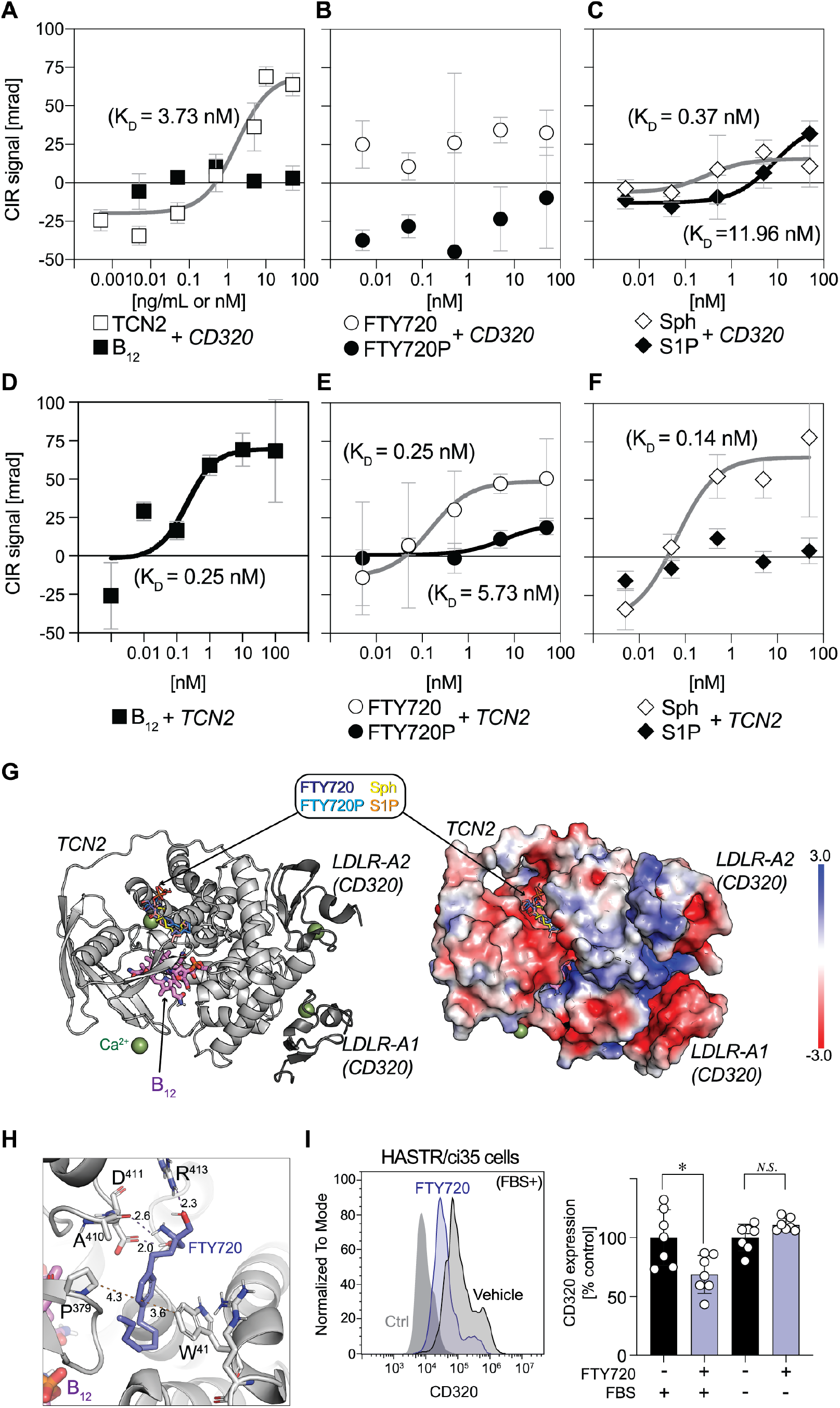
Direct binding of FTY720 with TCN2 induces astrocyte CD320 internalization. **A-C,** Specific binding curves between CD320 vs. TCN2 and B_12_ (**A**), FTY720 and FTY720P (**B**), sphingosine (Sph) and S1P (**C**). **D-F,** Specific binding curves between TCN2 vs. B_12_ (**D**), FTY720 and FTY720P (**E**), sphingosine (Sph) and S1P (**F**). Compensated Interferometric Reader (CIR) signals are plotted against concentrations of binding partners and fitted by nonlinear regression using the Michaelis-Menten equation (mean ± SEM, n = 6-7). **G,** Computational modeling of FTY720 binding site on TCN2 (PDB ID, 4ZRP). Cartoon representation of TCN2 (light grey) and LDLR domains of CD320 (dark grey) (*left*) and electrostatic surface of these complex (*right*). **H,** Magnified view of a potential FTY720 binding site in TCN2. Interactions are represented by dashed lines with distances between atoms (Å). **I,** A representative FCM histogram of CD320 expression on HASTR/ci35 cells stimulated with or without FTY720 (1 μM) in the presence or absence of FBS (*left*). Normalized CD320 expression levels (mean ± SEM, n = 6-7, pooled from 2 independent experiments). *, p < 0.05 by Kruskal-Wallis test with Dunn’s multiple comparisons test; N.S., non-significant.

Neither FTY720 nor Sph altered the affinity of TCN2 binding to B_12_ (**Fig. S4**), indicating distinct FTY720 binding sites compared to B_12_. Molecular modeling identified a potential binding site of FTY720 and Sph on the surface of TCN2, which was distinct from a B_12_ binding pocket (**Fig. 4G**). The model identified interactions between the benzene ring of FTY720 and W^41^/P^379^ of TCN2, and between a polar group of FTY720 and D^411^/R^413^ of TCN2 (**Fig. 4H**). Reduced binding of FTY720P or S1P to TCN2 might be explained by electrostatic repulsion between the phosphate groups of the ligands vs. the negatively charged surface of the binding site (**Fig. 4G**). Putative interacting residues require structural and mutagenesis studies for definitive assignments that are beyond the scope of this study but could be pursued in the future. Overall, these data supported TCN2 as a carrier and possible chaperone for both FTY720 and Sph, which also binds and transports B_12_ into astrocytes in conjunction with CD320.

Copious amounts of TCN2 is secreted into the plasma by the vascular endothelium and carries the newly absorbed B_12_ to all organs (Quadros and Sequeira, 2013). To assess whether FTY720 induces CD320 internalization in the presence of serum, it was evaluated by flow cytometry using an immortalized human astrocyte cell line, HASTR/ci35 cells (Furihata et al., 2016). FTY720 exposure decreased CD320 cell surface expression only in the presence of bovine serum that is known to contain TCN2, wherein FTY720/Sph binding could occur through conserved amino acid residues between human and bovine sequences (Polak et al., 1979). Internalization was not observed in the absence of serum (**Fig. 4I**). Sph also induced CD320 internalization (**Fig. S4**), indicating that FTY720- or Sph-bound TCN2 increases astrocytic B_12_ availability by driving CD320 internalization.

## Discussion

A molecular link between B_12_ and MS was identified through a novel CNS mechanism involving astrocytes that potentially explains the controversial relationship between B_12_ deficiency and MS that has spanned decades without resolution (Miller et al., 2005). The linkage consisted of 1) *Cd320* down-regulation in *ieAstrocytes* and reduced CD320 in human MS lesions; 2) worsening of disease by both CD320 genetic deletion and B_12_ deficiency revealed by EAE; 3) detection of specific, physical binding between FTY720 and the B_12_ carrier protein TCN2, and 4) CD320 internalization in astrocytes enhanced by FTY720. FTY720-relevant S1P_1_ receptor loss combined with increased IFN-I in EAE/MS (Teijaro et al., 2016; Yester et al., 2015) implicates a neuroprotection model involving astrocyte B_12_ signaling that is accessed clinically by fingolimod (FTY720) through a direct CNS mechanism (**Fig. 5**).

**Fig. 5.**
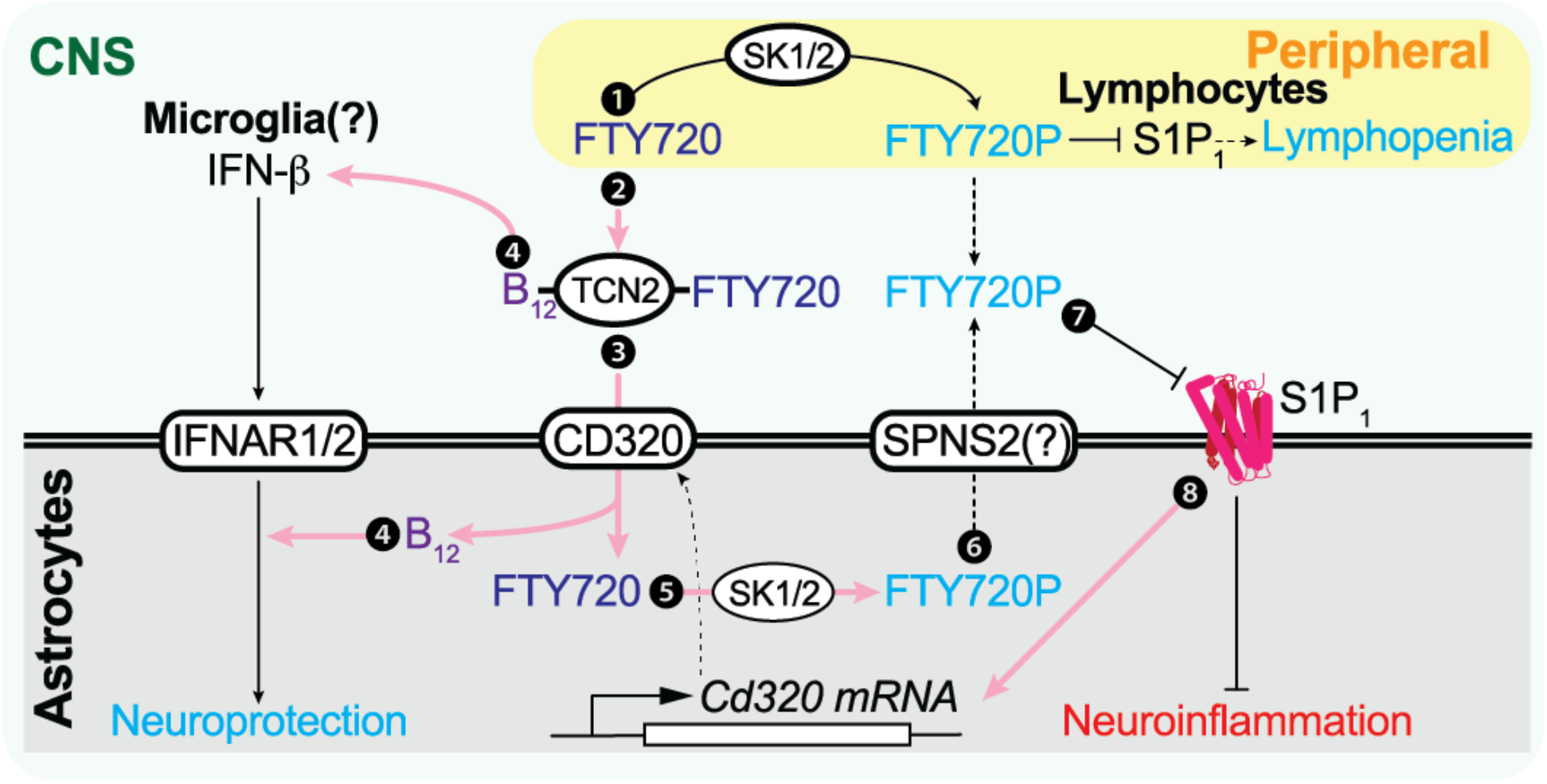
Schematic diagram of the FTY720 (fingolimod) CNS MOA through B_12_ and IFN-β signaling. Pink arrows indicate newly identified pathways that contribute to the direct CNS effects of FTY720 beyond functional antagonism of astrocyte S1P_1_. 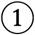; S1P_1_ inhibition in peripheral lymphocytes is the originally proposed CNS MOA. 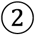; FTY720 is complexed with B_12_-TCN2. 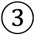; This FTY720 complex induces CD320 internalization and, along with B_12_, is taken up via astrocyte CD320. 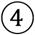; B_12_ is essential for astrocyte IFN-I sensitivity and microglial IFN-β production. 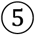; Astrocyte SK1/2 phosphorylates FTY720 to form FTY720P, which is identified in the accompanying Resource paper (Kihara, 2021). 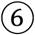; The resulting FTY720P may be transported via SPNS2 (Hisano et al., 2011) and functionally antagonizes S1P_1_ in an autocrine/paracrine manner. 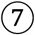; FTY720P functionally antagonizes astrocyte S1P_1_. 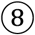; Astrocytic S1P_1_ inhibition increases CD320 expression, enabling increased engagement of, and astrocyte internalization, of the B_12_-TCN2-FTY720 complex.

### *CD320 is down-regulated in* ieAstrocytes *during neuroinflammation*

Astrocytes have long been considered bystanders in neuroinflammatory diseases (Rossi et al., 2007). However, identification of *ieAstrocytes* implicated the active involvement of astrocyte subpopulations in neuroinflammation that tracks with EAE disease severity (Groves et al., 2018). Molecular neuroinflammatory responses in *ieAstrocytes* are partly initiated by S1P-S1P_1_ signaling (Groves et al., 2018), resulting in c-Fos-dependent CD320 down-regulation (Jiang et al., 2010). Astrocytic CD320 down-regulation in both EAE SCs (**Fig. 1A-D**) and human MS plaques (**Fig. 1E-I**) are consistent with reported increases in S1P levels and S1P_1_ expression in both EAE SCs and human MS lesions (Kułakowska et al., 2010; Van Doorn et al., 2010). CD320 down-regulation could reduce astrocyte B_12_ availability that results in a negative feedback loop to down-regulate CD320 production (**Fig. S3**). CD320 expression in chronic EAE astrocytes maintained persistent down-regulation (Rothhammer et al., 2016), implicating *ieAstrocytes* as a potential precursor of reactive astrocytes. Another astrocyte classification nomenclature defined originally in mouse proposed “harmful” A1 and “helpful” A2 reactive astrocytes. An additional analysis of human MS astrocytes retained this nomenclature but utilized non-overlapping gene sets to define the A1 subtype (the A2 subtype was not reported in human brains) (Liddelow et al., 2017). *ieAstrocytes* and B_12_^free^ astrocytes expressed most of the proposed mouse A1/A2/Pan genes, but they could not be specifically categorized as A1- or A2-subtypes (**Fig. S1, S2**). It is possible that *ieAstrocytes* and related subtypes temporally precede A1/A2 subtypes. Developing an understanding of MS astrocyte subtypes warrant more investigation; however, our findings implicate *ieAstrocytes* defined by c-Fos expression as a key element of EAE/MS-specific reactive astrocytes.

### TCN2 functions as a sphingosine and FTY720 chaperone promoting CD320 signaling

In blood, S1P exists as a complex with its protein carrier chaperones that include albumin, apolipoprotein M (ApoM), and ApoA4, which presents S1P to its cognate receptors (Blaho, 2020). S1P concentrations are higher in blood than lymph, while sphingosine levels show an inverse trend (Nagahashi et al., 2016). TCN2 was identified here as a putative chaperone for sphingosine and FTY720. The Human Protein Atlas database reported the second highest expression level for TCN2 within lymphoid organs compared to all other organ systems (Colwill et al., 2011), which is consistent with higher sphingosine levels in the lymph (Nagahashi et al., 2016). Although neither FTY720 nor sphingosine altered the binding affinity of B_12_ to TCN2, both induced CD320 internalization (**Fig. 4 and Fig. S4**). Sphingosine and its chemical analogue FTY720 may thus act as TCN2 co-factors for the bound B_12_, forming a complex with CD320, thus enabling more B_12_ to enter astrocytes. Increased sphingosine levels in MS brains and EAE SCs (Miller et al., 2017) may reflect a compensatory mechanism to increase B_12_ in the brain (**Fig. 1**). The FTY720-TCN2 complex may also affect methylation processes of myelin proteins in oligodendrocytes via B_12_ signaling, which could be linked to the well-documented reversible demyelination in B_12_ deficiency (Miller et al., 2005). It is notable that B_12_ availability in the CNS through TCN2-CD320 may not be accurately reported by peripherally measured B_12_ levels which highlights a need to assess B_12_ brain levels.

### A direct CNS MOA for FTY720 is supported by astrocyte B_12_ and S1P_1_ signaling

Direct CNS activities of FTY720 beyond immunological mechanisms have received previous support from studies on S1P_1_-AsCKO mice challenged with EAE (Choi et al., 2011; Groves et al., 2018). These mice showed a loss of FTY720 efficacy along with a baseline reduction in clinical score despite maintaining FTY720’s immune cell trafficking effects (Choi et al., 2011). Complementary results were obtained here in EAE mice lacking either CD320 or B_12_ and which were also refractory to FTY720 therapy (**Figs. 1L and 2A**), but which showed a baseline *increase* in clinical disease score consistent with reduced CD320 gene expression from *ieAstrocytes* (**Fig. 1D**). FTY720 treatment that produces functional S1P_1_ antagonism also restored CD320 expression (**Fig. 1D**). This process requires SK1/2 activity that converts FTY720 to the active compound, FTY720P. In the accompanying paper (Kihara, 2021), single nucleus RNA-seq identified loss of *SPHK1/2*-expressing cells in secondary progressive MS (SPMS) brains. Further analyses using SK1/2-AsCKO mice showed a loss of FTY720 efficacy (Kihara, 2021). In this light, a failed Phase 3 clinical trial investigating FTY720’s efficacy in primary progressive MS (PPMS) patients (Lublin et al., 2016) may be explained by the loss of SKs from the PPMS brain.

These results, along with the expression of brain S1P receptors and accumulation of FTY720 in the CNS (Chun, 1999; Chun et al., 2000; Foster et al., 2007; Ishii et al., 2001; McGiffert et al., 2002; Zhang et al., 1999), implicate a direct CNS MOA for FTY720 that involves astrocytes (**Fig. 5**) that is amplified by B_12_ signaling. These direct CNS effects may also explain clinical data on reduced atrophy (De Stefano et al., 2017; Kappos et al., 2006; Yousuf et al., 2017) that contrasts with increased brain atrophy – “pseudoatrophy” – observed with the anti-inflammatory immunosuppressive agent natalizumab (Vidal-Jordana et al., 2013). In view of the current FTY720-B_12_ results, clinical reduction in brain atrophy accessed by fingolimod (FTY720) are consistent with increased brain atrophy reported under conditions of reduced B_12_, including that reported in CD320-KO mice (Arora et al., 2017) and human B_12_-deficient patients (Deng et al., 2017).

### Astrocyte IFN-I components are accessed by FTY720

IFN-I members include IFN-β, which was the first FDA-approved and current therapy for MS patients (IFN-β1b, Betaseron®)(Oh and O’Connor, 2015). It was developed as a peripheral immunomodulatory agent, while its CNS effects emerged later (Lublin, 2005). More severe EAE in IFN-β-KO mice (Teige et al., 2003) likely reflects the normally protective effects of CNS IFN-β production (Khorooshi et al., 2015; Kocur et al., 2015), a mechanism further supported by silencing an astrocytic IFN-I receptor (IFNAR1/*Ifnar1*), which aggravates EAE (Rothhammer et al., 2016). A new relationship between B_12_-TCN2-CD320-FTY720 and IFN-I signaling was identified here based on B_12_-deficient conditions and their effect on removing FTY720 efficacy, reducing astrocytic IFN-I sensitivity, and reducing endogenous IFN-β production (**Fig. 2**). A prior study reporting the positive effects of B_12_ on IFN-β therapy in EAE may reflect these CNS mechanisms (Mastronardi et al., 2004).

An additional relationship between S1P_1_ signaling and IFN-I was previously reported, where S1P_1_ activation inhibited IFN-I auto-amplification via c-Fos activation in astrocytes (Yester et al., 2015) while IFNAR1 degradation occurred in myeloid cells (Teijaro et al., 2016). S1P_1_ functional antagonism would counter these effects and it is consistent with IFN-β induction by FTY720 in EAE mice (**Fig. 3**). Notably, reduced IFN-β expression was not rescued by FTY720 in EAE mice generated with CD320 loss or B_12_^def^ (**Fig. 3**), underscoring the connected relationships amongst these elements (**Fig. 5**).

### Conclusions

A tangible molecular link between B_12_ deficiency and MS was identified through the effects on EAE of loss of B_12_ or CD320 in astrocytes, along with physical binding of B_12_-TCN2 with FTY720. B_12_ signaling appears to be capable of impacting fingolimod efficacy as well as disease severity mediated by sphingolipids in MS. The newly identified TCN2 binding of FTY720 promotes B_12_ and FTY720 availability within astrocytes. These data support the use of CNS penetrant B_12_ supplementation during fingolimod clinical treatment and suggest a need for assessing B_12_ signaling in the CNS beyond peripheral B_12_ measurements. Other S1P receptor modulators (Chun et al., 2021; Chun et al., 2019) may access one or more FTY720-B_12_ pathways without a requirement for phosphorylation by SK activity. Beyond MS (Noseworthy et al., 2000), neuroinflammation in the CNS has been associated with other brain diseases including, for example, Alzheimer’s disease (Acosta et al., 2017) and neuropsychiatric disorders (Trepanier et al., 2016), extending the potential biological and therapeutic relevance of astrocyte-B_12_ signaling.

## Acknowledgments

We thank Brinkmann, V. and Novartis for gifts of FTY720 and discussions. Wong, J., Mirendil, H., Blaho, V.A., Lee, M.H., Kachi, M., and all the lab members for discussions and support, and Jones, D. for editorial assistance. This work was supported by the National Institute of Neurological Disorders and Stroke of the National Institutes of Health under award number R01NS103940 (Y.K.), Novartis (J.C.), MEXT/JSPS KAKENHI 18H02627, 19KK0199, and 21H04798 (T.Y.), and 18K16246 and 21K08565 (H-C.L). Y.K. was supported by fellowships from the Uehara Memorial Foundation, Kanae Foundation for the Promotion of Medical Science, Mochida Memorial Foundation for Medical and Pharmaceutical Research, and the Human Frontier Science Program. A.G was supported by Medical Scientist Training Program and Pharmacology Training Grant at University of California San Diego (T32GM007752). The content is solely the responsibility of the authors and does not necessarily represent the official views of the National Institutes of Health.

## Author Contribution

Y.K. and J.C. conceived, designed, and supervised the project. D.J., Y.K., A.G., M.R., and R.R. performed *in vitro* and *in vivo* experiments and analyzed data; A.S., performed TCN2 modeling; H-C.L. quantified B_12_ levels under T.Y.’s supervision; T.F. provided HASTR/ci35 cells; E.V.Q. provided CD320-KO mice; Y.K., D.J., and J.C. wrote the manuscript. All authors contributed towards the final version of the manuscript.

## Materials and Methods

### Mice

All animal protocols were approved by the Institutional Animal Care and Use Committee (IACUC) of the Sanford Burnham Prebys Medical Discovery Institute and conform to National Institutes of Health guidelines and public law. The TetTag c-Fos reporter mice were generated by crossing Tg(Fos-tTA)1Mmay and Tg(tetO-HIST1H2BJ/GFP)47Efu/J mice, which expressed a green fluorescent protein-histone H2B fusion protein (GFP-H2B) under a tetO promoter controlled by a c-Fos inducible tetracycline transactivator (tTA) protein as described earlier (Tayler et al., 2011). S1P_1_^flox/flox^:hGFAP-cre mice (Choi et al., 2011) were crossed with TetTag mice for multiple generations to generate astrocyte specific S1P_1_ deletion (S1P_1_-AsCKO^fos^, TetTag:S1P_1_^flox/flox^:hGFAP-cre) and their littermate controls (WT^fos^, TetTag:S1P_1_^flox/flox^). TetTag mice were maintained on doxycycline chow (DOX, 40 mg/kg; Bio-Serv) throughout breeding, birth, and development to prevent GFP-H2B expression until experimental examination. CD320-KO mice were generated as described previously (Lai et al., 2013). C57BLJ/6 mice were maintained on a vitamin B_12_ deficient diet (Research Diets, Inc.) as necessary.

### EAE

EAE was induced in 7- to 13-wk old female mice as described previously (Kihara et al., 2009). Briefly, mice were immunized subcutaneously with 150 μg MOG35-55 (MEVGWYRSPFSRVVHLYRNGK, EZBiolab) in PBS and complete Freund’s adjuvant (BD Biosciences, cat # 263910) containing 4 mg/mL M. Tuberculosis H37Ra (BD Biosciences, cat # 231141) with or without intraperitoneal injection of 250 ng pertussis toxin (List Biological Laboratories, cat # 180) on day 0 and day 2. Daily clinical scores corresponding to the most severe sign observed were given as follows: 0, no sign; 0.5, mild loss of tail tone; 1.0, complete loss of tail tone; 1.5, mildly impaired righting reflex; 2.0, abnormal gait and/or impaired righting reflex; 2.5, hind limb paresis; 3.0, hind limb paralysis; 3.5, hind limb paralysis with hind body paresis; 4.0, hind and fore limb paralysis; and 4.5, death or severity necessitating euthanasia. Treatment of mice was performed by gavaging FTY720 (1 mg/kg; Novartis) and/or vitamin B_12_ (15 mg/kg; Sigma), unless otherwise noted.

#### Histological analysis

##### Formalin-fixed paraffin-embedded (FFPE) sectioning

SCs were incubated overnight in 10% neutral buffered formalin (PROTOCOL, Thermo Fisher Scientific) at room temperature. They were then cut into 3 sections and embedded in warm 2% agar (BD) dissolved in water and were allowed to solidify on crushed ice. The solid block was stored in 70% EtOH (PHARMCO, cat # 241000140CSGL), washed with 95% absolute EtOH, 100% absolute EtOH:Xylene (1:1), Xylene, molten warm Paraffin (Tissue-Tek, cat # 4005), and embedded into paraffin blocks using a manual paraffin embedder (Sakura Tissue-Tek). Sections (10 μm) were cut using a microtome (Leica RM2155) and used for Kluver Berrara staining and immunostaining. Histological scoring was performed as previously described (Kihara et al., 2005). Antigen retrieval with Diva Decloaker (Biocare) was performed followed by blocking with species-appropriate serum. Sections were incubated with primary antibodies including rabbit anti-CD320 (1:25 dilution, Proteintech, cat # 10343-1-AP), goat anti-Lcn2 (1:500 dilution, R&D systems, cat # AF1857), rabbit anti-Hmox1 (1:500 dilution, ThermoFisher Scientific, cat # PA5-27338), and chicken anti-GFAP(1:1000 dilution, Neuromics, cat # CH22102), followed by incubation with secondary antibodies conjugated with Alexa Fluor 488 or Alexa Fluor 568 (1:2000 dilution, ThermoFisher, cat # A11008 or cat # A11041, respectively) and counterstaining with DAPI (1:10,000 dilution, Sigma, cat # D8417). Sections were visualized and images acquired on a Zeiss Apotome.2 (Zen 2 Blue Edition).

#### RNA-seq

##### Nuclear isolation

SCs were rapidly dissected and frozen in liquid nitrogen. Samples were equilibrated to 4°C and dounce homogenized in a nuclei extraction buffer made with 0.32 M sucrose/5 mM CaCl_2_/3 mM Mg(CH_3_COO)_2_/0.1 mM EDTA/20 mM Tris-HCl pH 8.0/0.1% Triton X-100 in DEPC-treated H_2_O. Homogenized samples were filtered (50μm) (Celltrics, Sysmex) and washed in DEPC-treated PBS containing 2 mM EDTA (PBSE-d). Nuclei were purified by centrifugation at 3250*g* for 12 min in an iso-osmotic iodixanol gradient made of a 35%, 10%, and 5% OptiPrep (Millipore Sigma, cat # D1556) in DEPC-treated H_2_O containing 20 mM tricine-KOH (pH 7.8), 25 mM KCl, and 30 mM MgCl_2_ (Graham, 2001). Nuclei were recovered in the 35%-10% interface, washed in PBSE-d, and immunolabeled with rabbit anti-NeuN antibody (1:400 dilution, MilliporeSigma, cat # MABN140) in 1% bovine serum albumin (BSA)/PBSE-d for 20 min. Samples were washed in PBSE-d and labeled with a goat anti-rabbit APC conjugated secondary antibody (1:500 dilution, ThermoFisher Scientific, cat # 31984) and DAPI (1:5000 dilution, Sigma, cat # D8417) in 1%BSA/PBSE-d for 10 min. Samples were washed and suspended in PBSE-d. Nuclear populations were analyzed and sorted on a BD FACSAria Fusion. Gating was performed as follows: (i) DAPI positive, (ii) size and granularity consistent with nuclei by forward-scatter area (FSC-A) and side-scatter area (SSC-A), and (iii) single nuclei selected by both FSC-A and forward scatter height (FSC-H) and SSC-A and side-scatter height (SSC-H). Analysis was performed on FlowJo (10.0.8r1).

##### RNA isolation

Populations of 15,000 DAPI^+^NeuN^-^GFP-H2B^+^ nuclei were directly sorted into an extraction buffer for RNA isolation of Picopure™ RNA isolation kit (ThermoFisher Scientific, cat # KIT0204). RNase free Deoxyribonuclease I (DNase I) (Qiagen, cat # 1080901) was used to remove genomic DNA. For astrocyte RNA-seq, RNA was extracted using an RNeasy mini kit (Qiagen, cat # 74104) (with DNase I treatment) as per manufacturer’s instructions. RNA concentration and quality was assessed using a 2100 Bioanalyzer (Agilent Technologies).

##### Library preparation

Between 1-3 ng of the nuclear RNA was used in the SMART-seq v4 Ultra Low Input RNA Kit for Sequencing (Takara, cat # 634888) according to the manufacturer’s protocols. Briefly, first-strand synthesis by poly(A)-priming is followed by template switching and extension by reverse transcription, allowing high fidelity amplification of full-length cDNA transcripts by long distance PCR. A sequencing library was produced using 0.75 ng of the amplified cDNA in the Nextera XT Library Preparation Kit (Illumina, cat # FC-131-1024). For astrocyte RNA-seq, 150 ng of RNA per sample was employed in the generation of libraries using the NEB Next Ultra Pure RNA library kit for Illumina (NEB, cat # E7770S) per manufacturer’s instructions.

###### Nuclear RNA-seq analysis

Libraries were sequenced using the Illumina NextSeq 500 sequencer with 100 base pair single-end reads. Spliced Transcripts Alignment to a Reference (STAR) was used to align to the *Mus musculus* genome (mm10). All reads that mapped to exons and introns were used in this analysis, as nuclear RNA was previously noted to contain higher levels of intronic mapping (~27%) and intron expression is highly correlated with transcriptional activity (Gaidatzis et al., 2015; Grindberg et al., 2013). Partek software (v6.6) was used for nuclear RNA-seq analysis. The CPM values for each group were averaged to analyze fold differences between S1P_1_-AsCKO^fos^ and WT^fos^+FTY720 vs. WT^fos^. RNA-seq and data processing were performed at the Sequencing Core of The Scripps Research Institute.

###### Astrocyte RNA-seq analysis

Libraries were sequenced using the Illumina NextSeq 500 sequencer with 100 base pair single-end reads. Salmon (Patro et al., 2017) was used for transcript quantification while Counts Per Million (CPM) values were calculated using tximport (Soneson et al., 2015). RNA-seq and data processing were performed at the Sequencing Core of The Scripps Research Institute.

###### Astrocyte culture

Cerebral hemispheres of postnatal day 0 C57BL/6J pups were harvested and the meninges were removed. They were placed in DMEM/F12 (ThermoFisher Scientific, cat # 11320033), penicillin streptomycin (ThermoFisher Scientific, cat # 10378016), 1 mg/mL DNase I (Millipore Sigma, cat # D5025), and 0.05% trypsin (Millipore Sigma, cat # T1005). After gentle dissociation the tissue was incubated for 30 min at 37°C. Trypsin in the medium was inactivated by increasing the volume with DMEM/F12 containing 10% FBS (Gemini, cat # 100-500), centrifuged at 700 rpm for 5 min, filtered through a 40 μm cell strainer (Millipore Sigma, cat # CLS431750). The cells were cultured in DMEM/F12 containing 10% FBS and grown until 80% confluency before passage. The purity of astrocytes with this method was estimated to be more than 95% by immunostaining for glial fibrillary acidic protein (GFAP). For RNA-seq, after first passage, primary astrocytes were allowed to grow for 2 weeks either in normal media (DMEM/F12) or custom synthesized B_12_^free^ DMEM/F12 media (AthenaES) containing PS and 0.5% fatty acid free BSA.

Human astrocytes, HASTR/ci35 (Furihata et al., 2016), were cultured in Astrocyte Medium (ThermoFisher Scientific, cat # A1261301) at 33°C, 5% CO_2_ incubator. Cells were plated onto 6 well plates at 5 × 10^5^ cells/well one day before the experiment. Cells were washed with DMEM and stimulated with FTY720 and sphingosine in the presence or absence of serum for 1 h. Cells were detached and sequentially stained with anti-CD320 antibody (1:100 dilution: Proteintech, cat # 10343-1-AP) as the primary antibody and Alexa Fluor 488-conjugated Goat Anti-Rabbit IgG (1:1000 dilution: ThermoFishre Scientific, cat #A-11008) as a secondary antibody. FCM analyses were conducted with Novocyte (Agilent).

###### Quantitative PCR

The Superscript II first strand cDNA synthesis kit (ThermoFisher Scientific, cat # 18064022) was used to synthesize cDNA per manufacturer’s instructions for RT-PCR. This cDNA was used for qPCR analysis using GoTaq qPCR master mix (Promega, cat # A6001) using a Biorad CFX 384. Gene expression was normalized to β-actin. The primer sets for the target genes used in this study are provided in **Table S4.**

###### SDS page and western blotting

SC samples were minced for a few seconds intermittently in homogenization buffer (50 mM Tris-HCl pH 8.0, 150 mM NaCl, 0.5% Triton x-100, 2 mM EDTA, 1x protease inhibitor cocktail, and 250 mM sucrose). The mixture was incubated on ice for 1 h and centrifuged at 8000*g* for 10 min at 4°C. The supernatant was collected for measuring protein concentration using a Bradford reagent (Biorad, cat # 5000001). Samples were denatured in NuPAGE® LDS sample buffer (ThermoFisher Scientific, cat # NP0007) and 250 mM β-mercaptoethanol (Millipore Sigma, cat # M6250) at 80°C for 15 min, and run on 4-12% SDS Nu-PAGE gradient gel (ThermoFisher Scientific, cat # NP0323BOX). The proteins were transferred (NuPAGE® transfer buffer; ThrermoFisher Scientific, cat # NP00061) onto a PVDF membrane, blocked in 5% non-fat milk followed by incubation with an anti-CD320 Ab (1:100 dilution, Protein Tech, cat # 10343-1-AP). After several washes, the membrane was incubated in secondary antibody (HRP conjugated anti-rabbit or anti-mouse IgG(H+L) (1:10,000 dilution, Cell Signaling, cat # 7074P2 or 32230, respectively). Subsequently the signal was developed using Super Signal West Femto substrate (ThermoFisher Scientific, cat # 34094) and visualized using the molecular imager ChemiDoc™ XRS+ imaging system (Bio-Rad).

###### Binding assay

The Compensated Interferometric Reader (CIR) was developed from the backscattering interferometry (BSI) instrument that was described in detail previously (Mizuno et al., 2019; Ray et al., 2020). Recombinant human CD320 (R&D systems, cat # 1557-CD-050) and human TCN2 (R&D systems, cat # 7895-TC-050) were mixed with FTY720 (Novartis), FTY720P (Novartis), sphingosine (Avanti Polar Lipids, cat # 860490P), and S1P (Avanti Polar Lipids, cat # 860492P) and applied to the CIR. The *specific* binding signal was calculated by subtracting the control signal followed by normalization to the signal without ligand. The obtained signal displayed as milliradian were fitted by non-linear regression using the Michaelis-Menten equation using Prism software (GraphPad).

#### B_12_ measurement

Spinal cord tissues were crushed with an SK mill (SK-100; Tokken, Chiba, Japan). B_12_ was extracted with methanol by probe sonication. Cyanocobalamin-[^13^C_7_] (IsoSciences, King of Prussia, PA) was added as an internal standard. The tissue debris was removed by centrifugation at 10,000 × g for 5 min and filtration with YMC Duo-filter (YMC Co., Ltd., Kyoto, Japan) with a pore size of 0.2 μm, and a portion of the methanol extract was injected onto a Xevo™ TQ-S micro triple quadrupole mass spectrometry system (Waters) equipped with an electrospray ionization (ESI) source. Liquid chromatography (LC) separation was performed on a ZIC-pHILIC column (5 μm, 2.1 mm × 100 mm; Millipore) coupled to a ZIC-pHILIC guard column (2.1 mm × 20 mm; Millipore). Mobile phase A was acetonitrile containing 0.1% formic acid, and mobile phase B was water containing 0.1% formic acid. The LC method consisted of a linear gradient from 95% A to 50% B over 12 min at 0.2 mL/min, a linear gradient to 80% B over 16 min at 0.1 mL/min, 80% B for 20 min at 0.1 mL/min, 95% A for 7 min at 0.1 mL/min, and 95% A for 5 min at 0.2 mL/min (60 min total run time). The column temperature was set at room temperature. The detection was performed in multiple reaction monitoring (MRM) mode. MRM transitions from [M + H]^2+^ to the fragment ions at *m/z* 147.1 and 154.1 were used for B_12_ and the internal standard, respectively. The collision energy was set to 40 eV. The ESI capillary voltage was set at 1.0 kV, and the sampling cone was set at 30 V. The source temperature was set at 150°C, desolvation temperature was set at 500°C, and desolvation gas flow was 1000 L/h. The cone gas flow was set at 50 L/h.

#### Docking model

AutoDock is an automated docking method to identify binding modes of ligands inside the binding pocket of a biomolecular receptor (Goodsell et al., 1996; Morris et al., 1996; Morris et al., 2009). The computational protocol of AutoDock has been reported multiple times. In consistent with “united atom model”, only heavy atoms and polar hydrogens are used while docking simulations. To represent the target receptor, AutoDock uses a regularly spaced 3D grid which is defined by the user. A simplified representation of the receptor is then used during docking simulation. AutoDock family offers several tools, however AutoDock Vina is the latest method from that family (Trott and Olson, 2010). AutoDock Vina uses a unique approach to identify ligand binding modes implementing automatic calculation of the grid maps. Simultaneously, it also achieved approximately two orders of magnitude speed-up compared to its earlier version (AutoDock 4) while significantly improving the accuracy of the binding mode predictions based on an extensive training set. In the current report, we used Autodock (Version 4.2.3) and Vina (Version 1.1.2). The 3D coordinate of the protein bound to B_12_ was obtained from RCSB (http://www.rcsb.org/pdb/). Refining the system structure was achieved through using AutoDockTools. The 3D coordinate of the protein kept rigid while performing the docking. A graphical-user-interface for AutoDock, AutoDockTools (ADT, version 1.5.6 rc3), was used for adding hydrogens and Gasteiger charges and to generate the molecules in AutoDock suitable formats (pdbqt). A default grid spacing parameter (0.375Å) was used. The size of the grid box was chosen to be 100 × 100 × 100 which is large enough to cover the entire receptor and explore all possible binding pockets. For generating the docking parameter file (DPF) and the grid parameter file (GPF), ADT was used. AutoDock (4.2 and up) offers four different search algorithms based on speed and accuracy: 1) Simulated Annealing 2) Genetic Algorithm 3) Local Search and 4) Lamarkian. It has been reported multiples times that Genetic Algorithm (GA) often achieves most reliable results and highly appropriate for systems with many degrees of freedom. Thus, GA was used in the current report.

#### Statistical analysis

Results are expressed as mean ± SEM. Data were analyzed statistically by means of unpaired t test, and ANOVA with indicated post hoc tests as appropriate, using GraphPad Prism software. Values of p < 0.05 were considered to be statistically significant.

#### Data availability

RNA-seq data that supports the findings of this study are provided as Supplemental Tables (Table S1 and S3). Fastaq files have been deposited into Gene Expression Omnibus (GEO, https://www.ncbi.nlm.nih.gov/geo/). The accession number is GSE99115.

## Supplemental Figures provided as Ykihara_EAE-B_12__sup.docx

- Supplemental Figure S1. snRNA-seq and CD320 identification.
- Supplemental Figure S2. Analyses of B_12_^def^ EAE mice and B_12_^free^ astrocytes.
- Supplemental Figure S3. FTY720 efficacy in B_12_^def^ mice.
- Supplemental Figure S4. CIR-based binding assay and CD320 internalization.

## Supplemental Tables provided as YKihara_EAE-B12_SupTable.xlsx

- Supplemental Table S1. DEGs of nuclear RNA-seq, corresponding to Fig. 1C.
- Supplemental Table S2. Reactome pathway analysis of Cluster I genes that were commonly up-regulated in S1P_1_-AsCKO^fos^ and FTY720-treated groups as compared to WT^fos^
- Supplemental Table S3. DEGs of astrocyte RNA-seq, corresponding to Fig. 2D.
- Supplemental Table S4. Primer sets.

