## Supplemental Figures for "FTY720 requires vitamin B_12_-TCN2-CD320 signaling in astrocytes to reduce disease in an animal model of multiple sclerosis"

This file contains the following supplementary figures.

**Fig. S1.** snRNA-seq and CD320 identification.

**Fig. S2.** Analyses of $B_{12}^{def}$ EAE mice and $B_{12}^{free}$ astrocytes.

**Fig. S3.** FTY720 efficacy in $B_{12}^{def}$ mice.

**Fig. S4.** CIR-based binding assay and CD320 internalization.

***Table S1.** DEGs of nuclear RNA-seq, corresponding to Fig. 1C

***Table S2.** Reactome pathway analysis(Fabregat et al., 2017) of Cluster I genes that were commonly up-regulated in $S1P_1$-AsCKO$^{fos}$ and FTY720-treated groups as compared to WT$^{fos}$.

***Table S3.** DEGs of astrocyte RNA-seq, corresponding to Fig. 2D.

***Table S4.** Primer sets.

*Supplementaly tables are provided in an Excel spreadsheet in a different file.

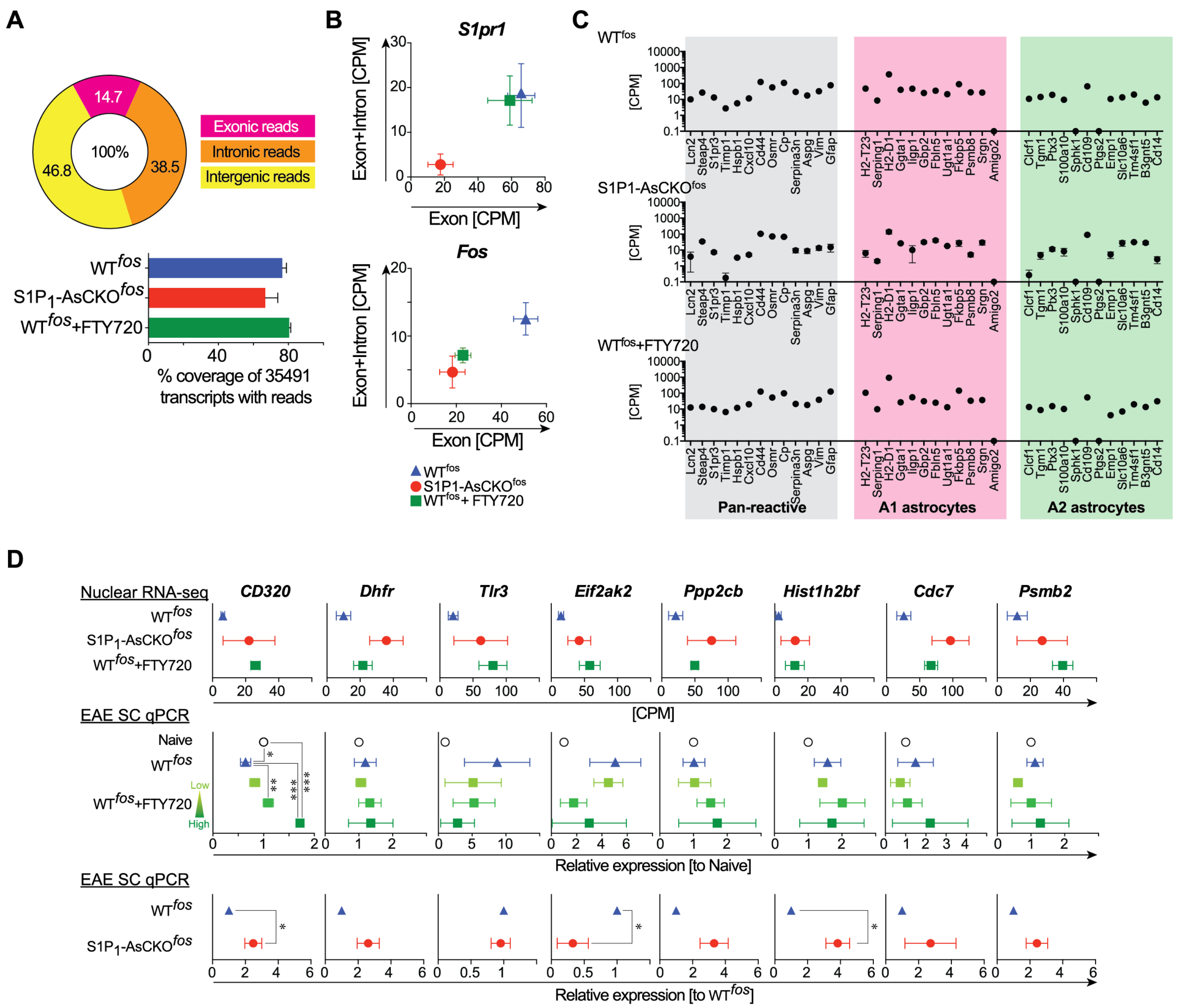


**E**

| | CD320-WT | CD320-KO |
|---|---|---|
| **Incidence** | 80% (20/25) | 89% (17/19) |
| **Mean day of onset** | 13.25 ± 0.53 | 11.24 ± 0.46 |
| **Mean MAX score** | 2.28 ± 0.24 | 2.63 ± 0.22 |
| **Cumulative score** | 15.9 ± 2.14 | 21.4 ± 1.98 |

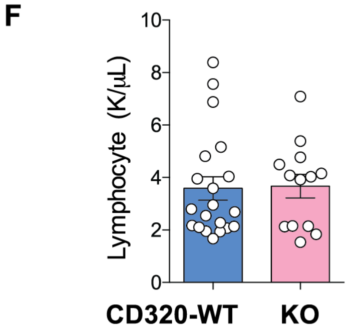


**Fig. S1. snRNA-seq and CD320 identification.**

**A,** averaged percentage of exonic, intronic, and intergenic reads in nuclear RNA-seq. The percent of reads that mapped to exons and introns were uniquely characteristic of nuclear RNA, with an increased representation of intronic reads. Percent coverage of transcripts (mean ± SEM). **B**, the exonic and exonic+intronic reads of

*S1pr1* were diminished only in $S1P_1$-AsCKO$^{fos}$ when compared to WT$^{fos}$ and WT$^{fos}$+FTY720, indicating that the DAPI$^+$NeuN$^-$GFP$^+$ nuclei were derived from astrocytes. The exonic and exonic+intronic reads of *fos* were diminished in both $S1P_1$-AsCKO$^{fos}$ and WT$^{fos}$+FTY720, indicating less astrocytic activation in these mice. **C**, comparison between nuclear RNA-seq data vs. recently proposed Pan/A1/A2 reactive astrocyte-specific genes. Although there was no obvious A1/A2-skewing (Liddelow et al., 2017) in DAPI$^+$NeuN$^-$GFP$^+$ populations, more than 90% of reactive astrocyte-specific genes were detected. **D**, expression profile of 8 genes identified in the pathway analysis that correspond to Fig. 1 C (mean ± SEM, *, $p < 0.05$; **, $p < 0.01$; *** $p < 0.001$ by one-way ANOVA in FTY720 groups, and one sample t test in KO groups). **E**, clinical parameters of EAE in CD320-WT and CD320-KO mice. Composite data from three independent experiments are shown. Cumulative scores were the sum of clinical scores from days 0 to 20 (means ± SEM, * $p < 0.05$ by Mann-Whitney U test). **F**, peripheral lymphocyte numbers. Each circle represents a single animal (mean ± SEM).

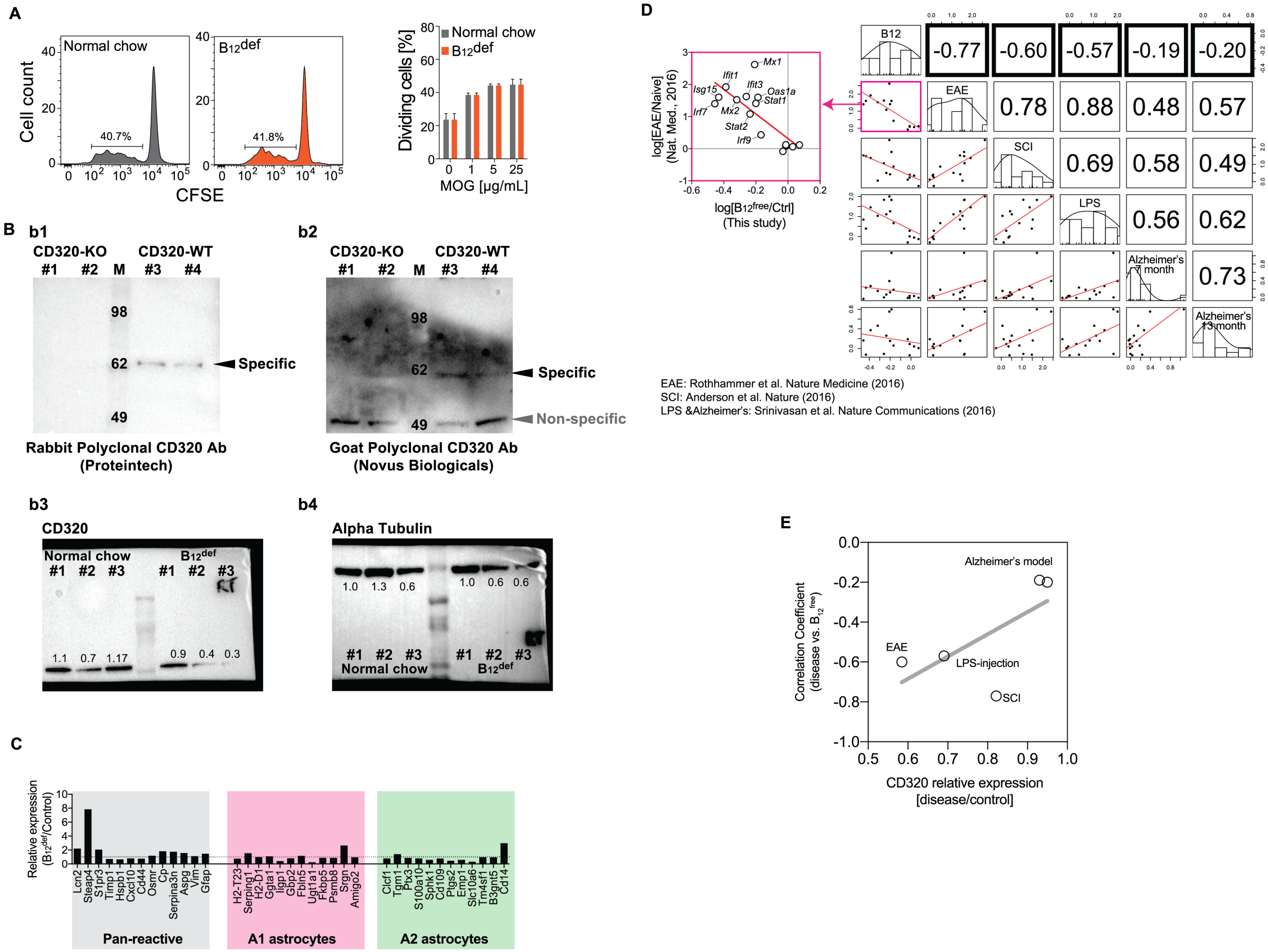

**Fig. S2. Analyses of $B_{12}^{def}$ EAE mice and $B_{12}^{free}$ astrocytes.**

**A**, T cell proliferation. Mononuclear cells were isolated from spleens of MOG35-55 immunized mice, and cultured in the presence or absence of $MOG_{35-55}$ for three days. Representative histograms of caboxyfluorescein succinimidyl ester (CFSE) intensities of $CD3^+$ T cells are shown from two independent experiments. Percentage of dividing T cells is indicated. MOG-induced cell proliferation is comparable in $B_{12}^{def}$ vs. controls (mean ± SEM, n = 3). **B**, Western blotting for CD320. (b1 and b2) Antibody specificities were assessed using proteins extracted from CD320-WT and CD320-KO SCs. The specific bands observed in CD320-WT disappeared in CD320-KO, indicating that both antibodies showed high specificity against mouse CD320. The calculated mass of CD320 is 27.7 kDa, but it is a heavily glycosylated protein (Quadros and Sequeira, 2013). (b3 and b4) CD320 expression was suppressed in EAE-induced $B_{12}^{def}$ mice as compared with controls. Numbers indicate relative intensities. **C**, Comparison between astrocyte RNA-seq data vs. recently proposed Pan/A1/A2 reactive astrocyte specific genes. No obvious A1/A2-skewing (Liddelow et al., 2017) was observed in $B_{12}^{def}$ astrocytes. **D**, Correlation matrix. Numbers indicate correlation coefficients. Correlation analyses of astrocytic IFN-I gene expression changes showed negative correlations between $B_{12}^{free}$ vs. EAE (Rothhammer et al., 2016), SC injury (Anderson et al., 2016), and LPS-injected mice (Srinivasan et al., 2016), but no correlation was found between $B_{12}^{free}$ vs. Alzheimer's model mice (Srinivasan et al., 2016). **E**, correlation of CD320 fold changes vs. correlation coefficients between $B_{12}^{free}$ (bold boxes in D) and indicated disease conditions. Moreover, these correlation coefficients were positively correlated with the extent of astrocytic Cd320 down-regulation in these diseases, suggesting an association of astrocytic IFN-I sensitivity with the $B_{12}$-CD320 pathway in neuroinflammation.

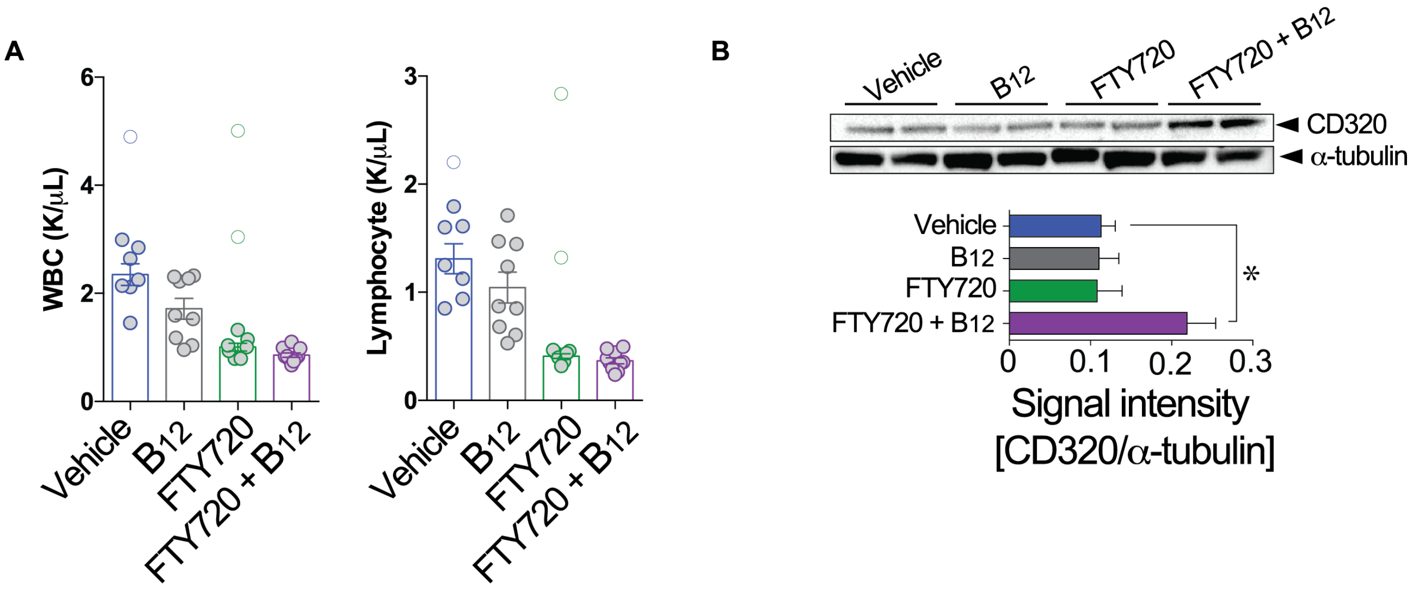


**Fig. S3. FTY720 efficacy in $B_{12}^{def}$ mice.**

**A**, peripheral white blood cell (WBC) and lymphocyte counts in EAE mice treated with vehicle, $B_{12}$, FTY720, and FTY720+$B_{12}$. Lymphocyte trafficking effects of FTY720 appear to be intact for all experimental conditions examined. Each point represents a single animal. Some samples (open circles) were eliminated from the datasets based on Smirnov-Grubbs test before performing statistical analysis by Kruskal-Wallis test with Dunn's multiple comparisons test (mean ± SEM). **B**, Western blotting for CD320 in SCs of EAE mice at 50 days post immunization and 25 days post treatment. Signal intensity of CD320 was normalized by comparison to α-tubulin (mean ± SEM, n = 4 animals, * $p < 0.05$ by one-way ANOVA with Bonferroni's multiple comparisons test).

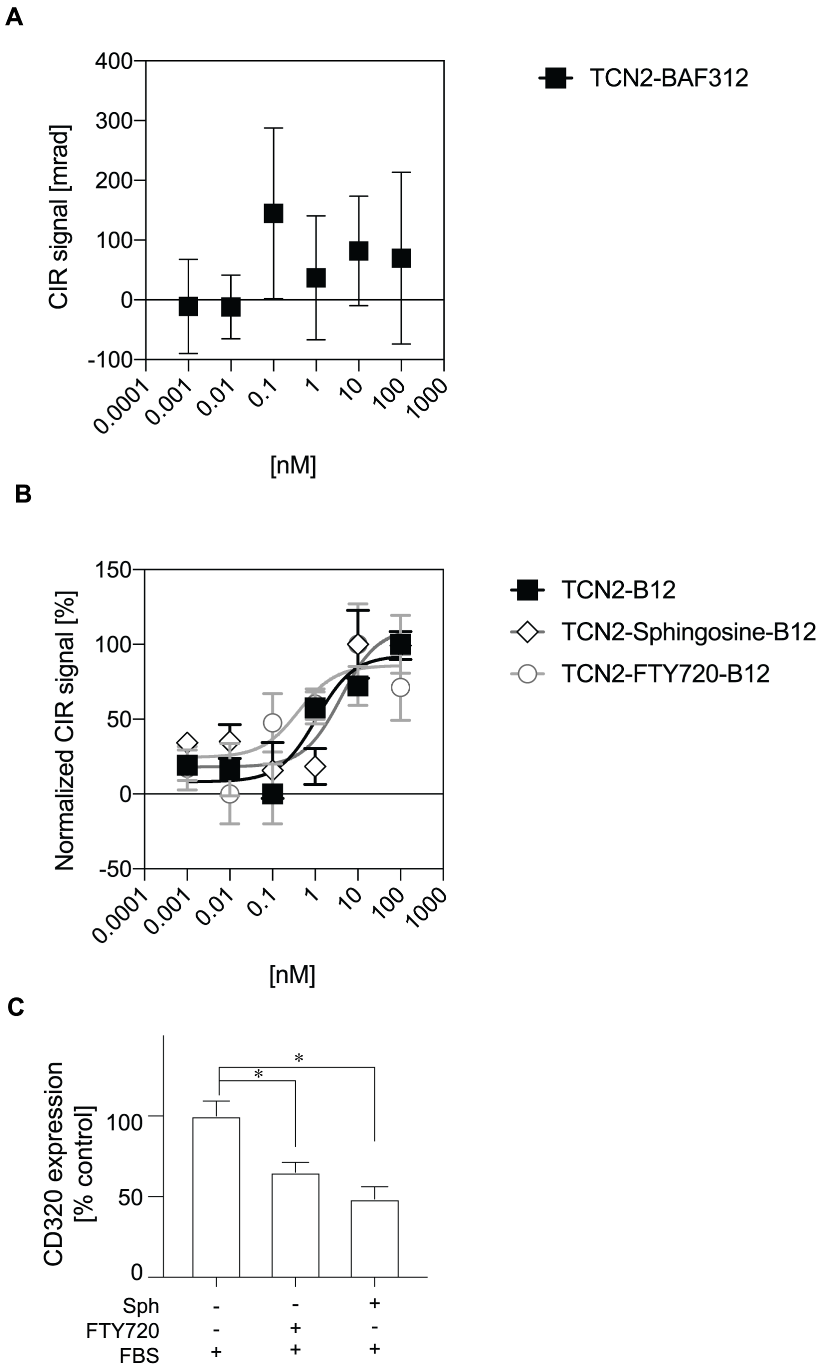


**Fig. S4. CIR-based binding assay and CD320 internalization.** **A**, No clear binding signals between TCN2 vs. BAF312 (siponimod). **B**, Specific binding curves between TCN2-$B_{12}$ vs. FTY720 and sphingosine. Compensated Interferometric Reader (CIR) signals are plotted against concentrations of binding partners, normalized, and fitted by nonlinear regression using the Michaelis-Menten equation (mean ± SEM, n = 4-5). **C**, CD320 internalization in HASTR/ci37 cells. Cells were stimulated with 1 µM FTY720 or 1 µM Sph in the presence of FBS. * $p < 0.05$, one-way ANOVA with Bonferroni's multiple comparisons test.
